# The RLR-MAVS-IRF3 axis activates the IFN pathway to restrict Tonate virus (TONV) infection

**DOI:** 10.64898/2026.07.09.737427

**Authors:** Thomas Labadie, Roger J. Eloiflin, Zoé Denis, Marie Motos, Joanna Re, Moritz Schussler, Morgane Chemarin, Isabelle Moltini-Conclois, Valerie Courgnaud, Dorothée Missé, Nadine Laguette, Karim Majzoub

## Abstract

Tonate virus (TONV) is a neglected mosquito-borne alphavirus of the Venezuelan equine encephalitis complex associated with febrile illness, encephalitis, and fetal central nervous system abnormalities. Yet, host pathways that sense TONV infection and restrict its replication remain poorly defined. Here, we investigated the interaction between TONV and the type I interferon (IFN-I) system in human cells. We show that TONV infection led to the accumulation of cytosolic double-stranded RNA and a robust IFN response. RIG-I and MDA5 depletion as well as that of MAVS and IRF3 strongly reduced TONV-induced IFN response. Disruption of RIG-I, MDA5, MAVS, or IRF3 resulted in an increase of dsRNA-positive cells and a higher viral RNA accumulation. Finally, we found that treatment with exogenous IFN-I strongly inhibits TONV replication reducing both viral RNA loads and infectious particle production. Thus, together, our results identify the RIG-I/MDA5-MAVS-IRF3 axis as a major pathway sensing TONV infection and establishing an IFN-I-dependent antiviral state, providing the first molecular characterization of TONV innate immune sensing in human cells.

## Introduction

Tonate virus (TONV) is a neglected mosquito-borne alphavirus for which surprisingly little is known despite decades of circulation in South America. TONV has been historically classified as belonging to the Venezuelan equine encephalitis (VEE) virus subtype IIIB complex ^1,2^, placing it among the New World encephalitic alphaviruses that include Venezuelan equine encephalitis virus (VEEV) and related viruses with recognized neurotropism and emergence potential ^3–5^. Across the alphavirus genus, and particularly within the VEE complex, the outcome of infection is strongly shaped by the type I interferon (IFN-I) response: viral strains differ in their sensitivity to IFN-I ^6,7^, which correlates with attenuation and virulence in VEEV ^6,8^ and alphaviruses have evolved multiple mechanisms to counteract innate immune mechanisms ^9–11^. In parallel, alphavirus RNAs are recognized by cytosolic RNA sensors, and work on VEEV and sindbis virus (SINV) has shown that RIG-I-like receptors (RLRs) can drive MAVS- and IRF3-dependent antiviral programs ^12,13^, although the contribution of these pathways varies with the virus and the cellular context ^13,14^. Whether TONV follows this broader alphavirus paradigm, however, has remained unknown.

This lack of mechanistic insight into TONV biology is notable since available epidemiologic and clinical data suggest that TONV is mostly detected within a defined geographical region in the north of Latin America^1,5,15^. First identified in French Guiana in the 1970s^16^, TONV has since been associated with human exposure in endemic areas, with higher seroprevalence reported in some populations from savannah regions such as Bas Maroni, west of French Guiana ^17^. Most recognized infections appear to present as a nonspecific arboviral febrile illness, which likely contributes to under-diagnosis, especially in settings where dengue and other arboviruses co-circulate^3^. More broadly, endemic VEE complex infections have been often described as being hidden “under the dengue umbrella” because of their overlapping clinical manifestations and limited access to specific diagnostics ^17,18^. In some cases, severe disease can occur: fatal encephalitis and vertical transmission with severe fetal central nervous system lesions have both also been reported ^4,19^. The ecology of TONV also remains only partially defined, with isolations from mosquitoes, birds and nest bugs, and more recently bats, supporting persistent enzootic circulation rather than a fully understood transmission cycle ^5,20,21^. Importantly, recent experimental work showed that *Aedes albopictus* is competent for TONV transmission, a finding that warrants attention given the continuing habitat expansion of *Aedes* mosquitoes under climate change and the broader redistribution of *Aedes*-borne arboviral risk, even if the epidemic potential of TONV remains unproven to date ^22,23^.

Against this background of sparse knowledge, likely under-recognition, and documented neurologic and fetal complications, defining how TONV is sensed in human cells is important both for understanding antiviral restriction and for improving preparedness toward neglected alphaviruses with potential for emergence. We therefore investigated the interaction of TONV with the human IFN-I system, more specifically the upstream pathways responsible for innate immune viral sensing. We found, that similarly to other alphaviruses ^9^, TONV is both sensitive to IFN and capable of inducing IFN production through a RIG-I/MDA5-MAVS-IRF3-dependent pathway.

## Results

We first asked whether TONV infection elicits a type I IFN response in human cells. Because TONV has been described to display neuronal tropism, this was assessed on the T98G a fibroblast-*like* cell line derived from a human glioblastoma ^24^. Quantification of mRNA levels by RT-qPCR showed that *IFNβ* and Interferon-stimulated genes increased over the time course of infection and their levels correlate with the TONV virus in a MOI-dependent manner (Figure 1A-C, Figure S1A-B). Similar results were obtained in A549 cells derived from an epithelial human lung carcinoma ^25^ (Figure 1A-C). We then measured the secretion of bioactive type I IFN cytokines from both T98G and A549 cells infected with TONV using a reporter assay ^26^ and found that similar to mRNA levels, type I IFN cytokine production in infected cells is stimulated by TONV infection in a MOI-dependent manner (Figure 1D). This suggests that TONV infection leads to type I IFN responses susceptible of promoting control of viral infection.

**Figure 1.**
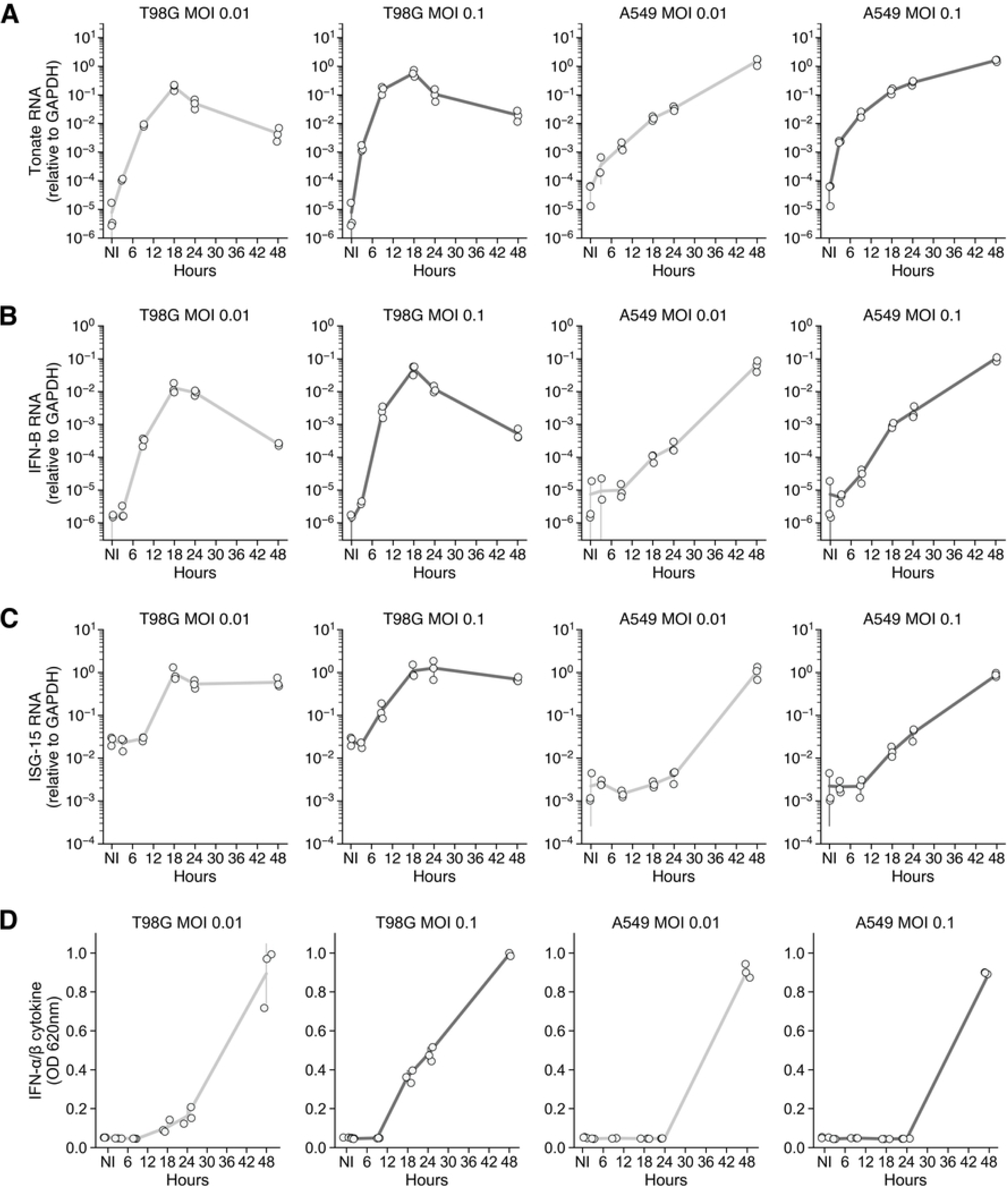

Thus, we next assessed whether TONV is sensitive to type I IFNs. To this end, we pre-treated A549 and T98G cells with increasing units of type I IFN-α cytokine 24 h prior to TONV infection. This treatment markedly decreased both viral RNA loads (Figure 2A) and infectious particle production (Figure 2B, Figure S2A), regardless of the Multiplicity of Infection (MOI) (Figure 2A, B). We also confirm that IFN pre-treatment increases ISGs levels in both used cell lines (Figure S2B). Our results show that, similar to other alphaviruses, TONV is restricted in cells stimulated with IFN, likely through the action of IFN-stimulated genes ^7,27,28^.

**Figure 2.**
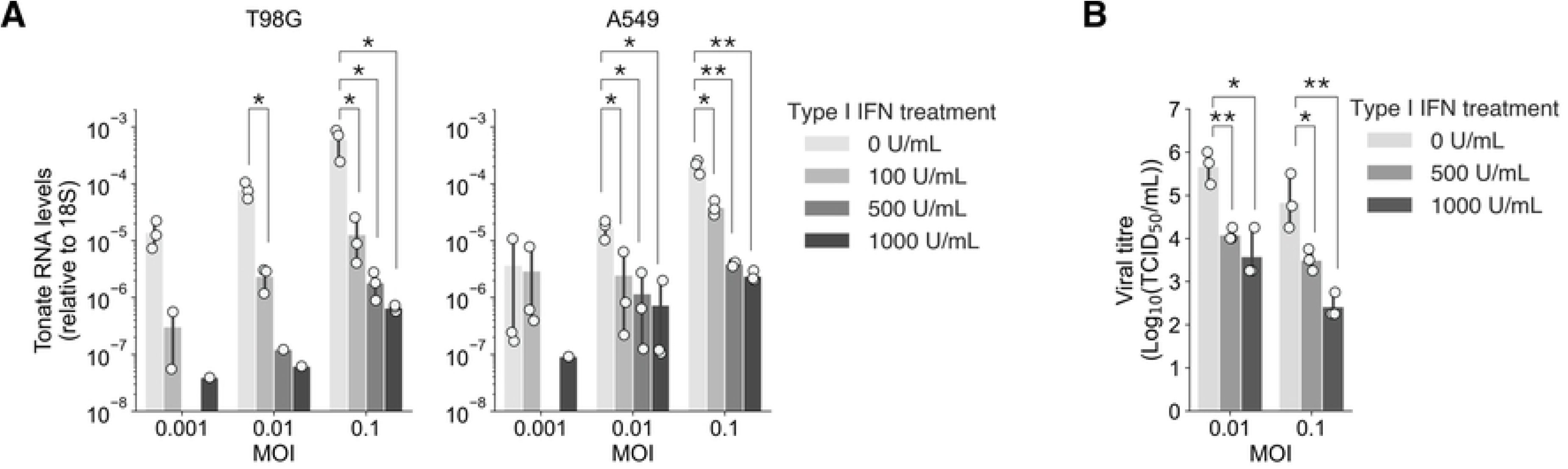

Our results show that TONV infection induces the IFN pathway (Figure 1) and its replication can be efficiently inhibited when the IFN I cytokine is added to cells exogenously before viral infection (Figure 2). To test whether the RLR-dependent viral sensing pathway may be involved in TONV detection, we generated knocked-out (K.O.) cells, using a CRISPR-Cas9 approach ^29^, of two previously described cytosolic viral RNA (vRNA) sensors: retinoic acid-inducible gene I (RIG-I) and melanoma differentiation-associated protein 5 (MDA5) (Figure 3A), both shown to detect vRNA of other alphaviruses (e.g. SINV, VEEV and Semliki Forest virus (SFV)) and to induce an antiviral state ^12,30^. Since RIG-I and MDA5 can have complementary or redundant functions, we also generated K.O. cells for both sensors (Figure 3A). We also generated K.O. cells for Mitochondrial antiviral-signaling protein (MAVS) and Interferon regulatory factor 3 (IRF3) (Figure 3A), two key players in the viral sensing signal transduction pathway, acting downstream of RLRs ^31^. Indeed, the products of these two genes are essential for IFN pathway induction. MAVS is a central adaptor protein downstream of RIG-I and MDA5 that relays cytosolic viral RNA sensing to the TBK1/IKKε kinase complex, leading to IRF3 activation. Phosphorylated IRF3 then translocates to the nucleus and drives transcription of IFN-β and other type I interferon genes^12,32–34^. We found that knocking out each of these genes resulted in higher TONV RNA loads, with the most pronounced effect observed in the absence of IRF3 (Figure 3B). As expected, *IFNβ* cytokine production was strongly impaired in MAVS and IRF3 KOs, indicating that MAVS-IRF3 axis controls IFN responses to TONV infection, with the most pronounced effect found in IRF3-K.O.s (Figure 3C). Interestingly, both MDA-5 and RIG-I KOs had lower *IFNβ* production compared to Wild-type (WT) cells upon TONV infection (Figure 3C) and the double RIG-I/MDA5 K.O. displays an additive effect, with *IFNβ* levels similar to those produced in the MAVS K.O. (Figure 3C). The impaired *IFNβ* production is also reflected by reduced ISG15 (Figure 3D) and Interferon-induced GTP-binding protein MX1 (Figure S3) mRNA levels in KO cells. These observations argue for a redundant role of MDA5 and RIG-I in TONV vRNA sensing and activation of the IFN pathway. Indeed, an earlier study have shown that both RIG-I and MDA5 can detect alphavirus replication, including during VEEV infection, in a concentration-dependent manner ^12^.

**Figure 3.**
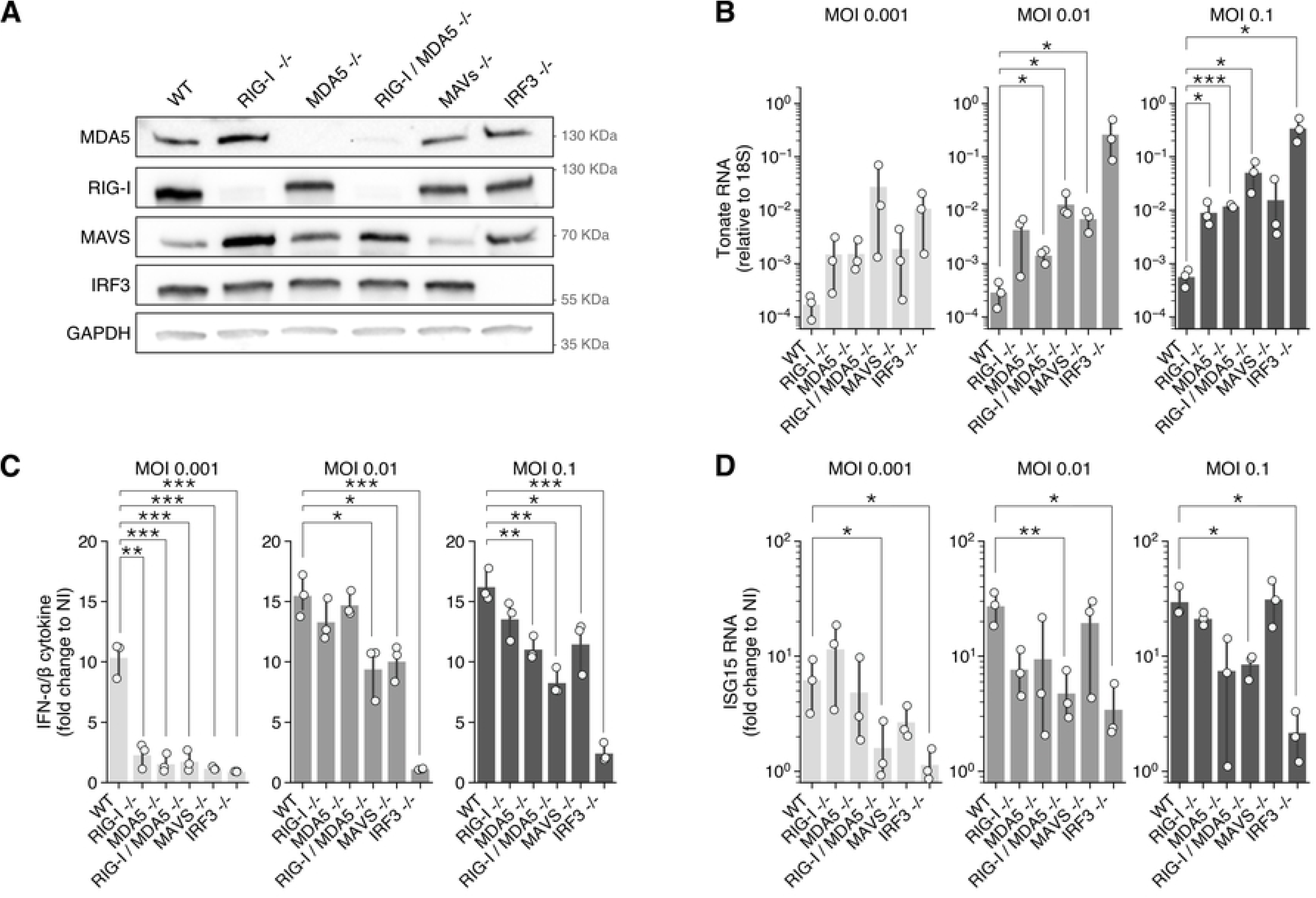

Next, we sought to determine if the impairment of TONV-induced IFN responses have any functional consequence on intracellular viral RNA accumulation. We first wanted to determine if TONV replication causes the formation of double stranded RNA (dsRNA), a known viral replication intermediate recognized by RLRs as a Pathogen Associated Molecular Pattern (PAMP) ^31^. To this end, T98G cells were infected with TONV at different MOIs for 24 h, fixed and immunostained with the J-2 antibody, an anti-dsRNA antibody widely used to detect replication-associated viral dsRNA intermediates in infected cells ^35^. Our results indicate that TONV infection induces the formation of cytosolic dsRNA, readily visible in infected cells by confocal microscopy (Figure 4A).

**Figure 4.**
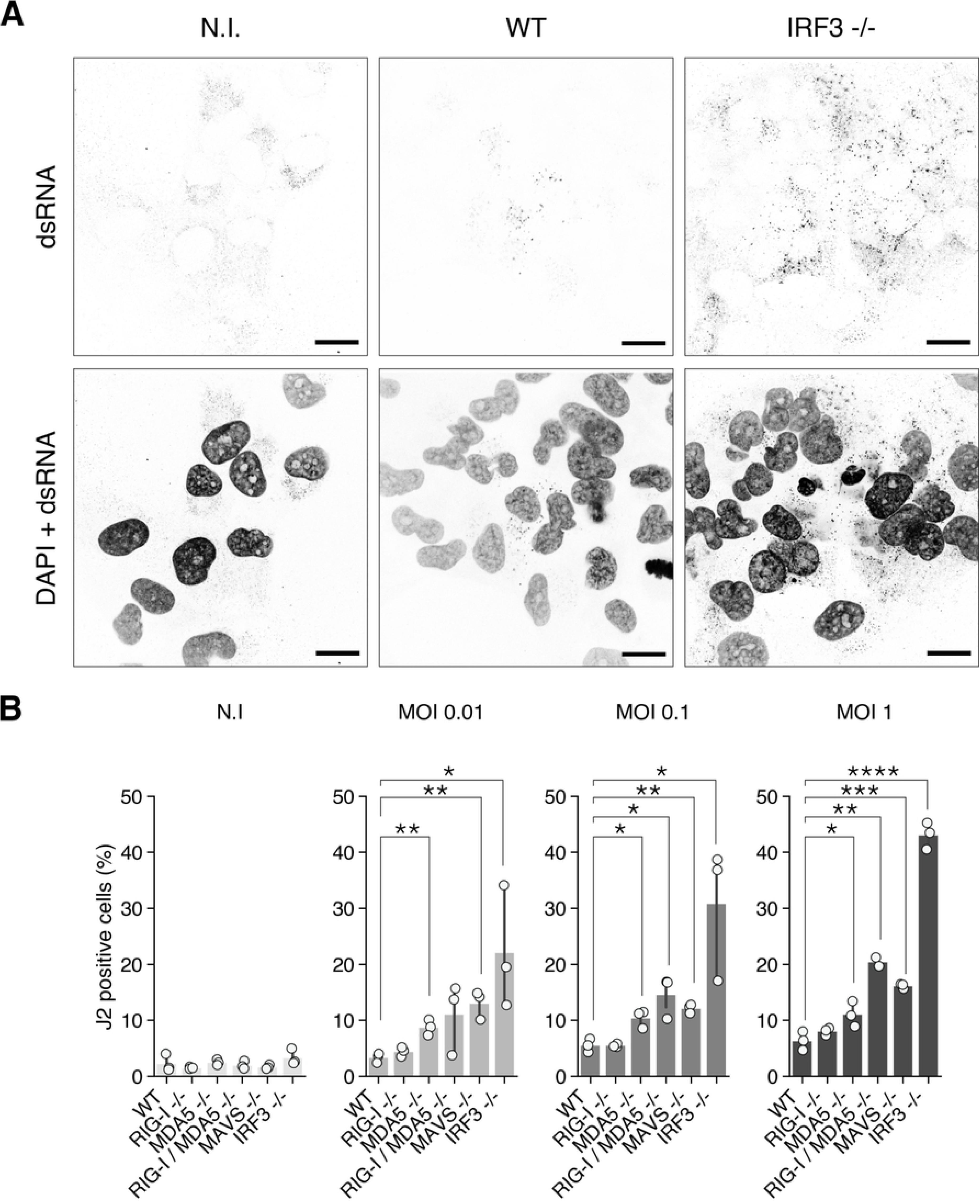

As dsRNA accumulation can be considered a surrogate for viral replication, we quantified the percentage of dsRNA-positive cells in RIG-I, MDA5, MAVS and IRF3 KO cells, compared to WT cells, to determine if the lack of these factors can affect viral replication (Figure 4B). All KO cell lines displayed higher J2-staining than WT cells, with IRF3 KOs showing the highest amounts of dsRNA accumulation, followed by MAVS KOs and then the RLR KOs (RIG-I and MDA5). These results are in agreement with viral RNA loads measured by by RT-qPCR (Figure 3B). In conclusion, our results suggest that TONV is sensed by the RLR-MAVS-IRF3 pathway, activating the IFN pathway to limit viral replication.

## Discussion

TONV remains one of the least characterized mosquito-borne alphaviruses despite evidence of long-term circulation in French Guiana and neighboring regions, documented human exposure, and reports of severe neurologic and fetal disease in rare cases ^3–5,17,19^. In this study, we show that TONV is strongly restricted by exogenous type I interferon (IFN-I) and, conversely, that infection itself induces an IFN response in human cells. Mechanistically, our data identify the RIG-I/MDA5-MAVS-IRF3 axis as a major pathway controlling this response and limiting viral replication. Thus, although TONV has remained largely neglected, our results now place this virus within a now clearer innate immune framework and show that its interaction with the host IFN system broadly resembles that of other alphaviruses ^9,12,30,34^.

A first important finding of this work is that TONV is highly sensitive to IFN pretreatment in two distinct human cell lines. This is consistent with earlier studies showing that alphavirus infection outcomes are strongly shaped by the type I IFN response, and that IFN sensitivity can correlate with attenuation and virulence, particularly within the VEE complex ^7,8,36^. While this may be expected based on TONV taxonomy, it remained important to demonstrate experimentally. Demonstrating IFN sensitivity provides a useful reference point for future work on TONV virulence, cell tropism, and possible immune evasion strategies.

Our data further show that TONV is not only restricted by IFN but is also capable of inducing the IFN pathway. In infected T98G cells, TONV induced IFNβ and ISG15 transcripts and triggered secretion of functionally active type I IFN cytokines, indicating that the virus is efficiently sensed by host cells. This fits well with the broader alphavirus picture, in which replication-associated RNA species generated in the cytoplasm activate innate immune sensing pathways and induce antiviral transcriptional programs ^9,12,30,34^. The detection of dsRNA in TONV-infected cells by J2 staining further supports this model, as dsRNA is a well-established viral PAMP generated during the replication of many positive-strand RNA viruses ^31,35,37^.

A central contribution of this study is the identification of the RLR-MAVS-IRF3 axis as the major pathway responsible for TONV-induced IFN production. Both RIG-I and MDA5 contributed to IFNβ induction, whereas deletion of both RLRs and of MAVS or IRF3 had a stronger effect on IFN production and was associated with increased viral accumulation. This pattern is in agreement with previous studies on alphaviruses showing that RIG-I and MDA5 can both detect replication-associated RNAs and act in a partially redundant or complementary manner depending on viral burden and cellular context ^12,30,38^. Our observation that single RIG-I or MDA5 knockout reduced *IFNβ* mRNA induction without markedly affecting total IFN bioactivity or ISG15 induction is consistent with this model. By contrast, the stronger phenotype observed in MAVS-and IRF3-deficient cells is expected for factors positioned downstream of both sensors and converging on the core pathway of IFN induction ^32–34^. Together, these findings support a model in which TONV-generated RNA species are sensed by both RIG-I and MDA5, then signal through MAVS to activate IRF3 and establish an antiviral state that restricts viral replication. Importantly, the use of T98G cells is also noteworthy in this context. Although our study was not designed as a neuropathogenesis model, TONV has been associated with encephalitis in humans and with severe fetal central nervous system abnormalities ^4,19^. Demonstrating robust IFN induction and innate sensing in a human brain-derived cell line does not only offer an interesting model to test antivirals, it also adds biological relevance to our findings and supports the idea that antiviral recognition of TONV may be particularly important in tissues linked to clinically severe disease ^4,19^.

Beyond the mechanistic findings, this work is relevant because TONV sits at the intersection of neglect, diagnostic uncertainty, and emergence concern. Clinically, TONV infection can resemble other arboviral febrile illnesses and may therefore remain underestimated in endemic areas where dengue and related arboviruses dominate diagnostic attention ^3,17,18^. At the same time, severe neurologic disease and vertical transmission have been documented ^4,19^, indicating that TONV infection is not trivial. Recent evidence that *Aedes albopictus* is competent to transmit TONV further increases the importance of understanding this virus before any broader expansion in transmission range ^22,23^. While the true epidemic potential of TONV remains unknown, our work provide key foundational molecular characterization that is critical in case of larger emergence and provides an immediate framework for future work on pathogenesis, immune control, and antiviral interventions.

## Limitations of the study

Our study has obviously limitations. It was performed in immortalized human cell lines, which are experimentally useful but do not capture the full diversity of relevant target cells *in vivo*. In addition, although our knockout approach identifies the major sensing pathway, it does not define the exact vRNA species recognized during infection or the precise relative contribution of RIG-I and MDA5 over time. Future studies should address these points and also determine whether TONV, like other alphaviruses, encodes specific antagonists of IFN induction or signaling^10,39–41^.

## Materials and Methods

### Cells and infection

Wild-type (WT) fibroblast-like cell line derived from a human glioblastoma T98G and WT epithelial human lung carcinoma A549 were a gift from Caroline Goujon (Institut de Recherche en Infectiologie de Montpellier, Montpellier, France). WT and genetically modified T98G and A549 cells were maintained in Dulbecco’s Modified Eagle Medium (DMEM) supplemented with 10 % Foetal Bovine Serum (FBS, Eurobio), 1% L-glutamine (Lonza), 1% Penicillin/Streptomycin (Lonza). In-house-generated knockout cell lines were maintained in the presence of the puromycin selection antibiotic. All cell lines were maintained at 37°C under 5% CO_2_. T98G and A549 cells were infected by inoculation of Tonate virus CaAn 410d strain (CRORA collection, Institut Pasteur, Paris) diluted in cell culture medium (supplemented with 10% FBS) at a multiplicity of infection (MOI) of 0.001, 0.01, 0.1 or 1 plaques forming unit per cell (pfu/cell). Cells were maintained for 24 h at 37°C before further processing for downstream analysis. When mentioned, cells were pre-treated with Universal Type I IFN (Human IFN-Alpha Hybrid Protein, PBL Assay Science) 18h after cells seeding in culture dishes. Cells were maintained for another 18h in presence of IFN before renewing the cell culture medium and infection with TONV.

### Plasmids, constructs, and synthetic nucleic acid probes

T98G^RIG-I-/-^, T98G^MDA5-I-/-^, T98G^RIG-I-/- MDA5-/-^, T98G^MAVS-/-^ and T98G^IRF3-/-^ cell lines were generated by lentiviral transduction followed by selection, using the LentiCRISPRv2puro plasmid (Addgene #52961) in which targeting guide RNAs (gRNA) from the Human CRISPR Knockout Library (GeCKO v2) were cloned using the following sequences (Table 1). Synthetic primers (gRNAs) were purchased from Integrated DNA Technologies (IDT) before cloning in PlentiCRISPRv2.

**Table 1.**
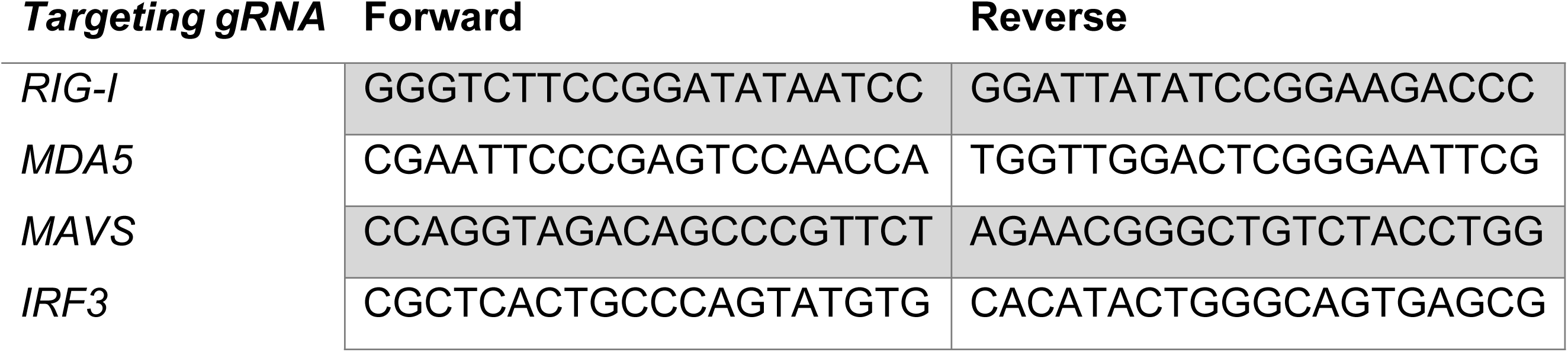
Targeting guide RNA (gRNA) used for lentiviral constructions.

### Generation of knock-out cell lines

Lentiviral particles used to generate T98G^RIG-I-/-^, T98G^MDA5-I-/-^, T98G^RIG-I-/- MDA5-/-^, T98G^MAVS-/-^ and T98G^IRF3-/-^ cell lines were produced by co-transfection of 2 × 106 HEK293T with 1 μg of LentiCRISPRv2puro plasmid expressing targeting gRNAs, together with 0.5 μg of psPAX2, and 0.5 μg of pMD2.G, using the calcium phosphate transfection protocol. Lentiviral particles were harvested 48 h after transfection and filtered (0.45 µM) prior to transduction of WT T98G. Selection was initiated 72 h post-transduction using 2 μg/ml puromycin. Protein levels of different target genes were controlled by Western blot (WB) after 6 days of selection.

### Whole-cell lysate preparation and WB

T98G WT and KO cells were lysed in five packed cell volumes of TENTG-150 (20 mM Tris-HCl [pH 7.4], 0.5 mM EDTA, 150 mM NaCl, 10 mM KCl, 0.5% Triton X-100, 1.5 mM MgCl2, and 10% glycerol, supplemented with 10 mM β-mercaptoethanol, 0.5 mM PMSF, and phosphatase inhibitor [Sigma-Aldrich]) for 30 min at 4°C. Lysates were centrifuged for 30 min at 14,000 g, and supernatants were collected for WB. Protein quantification was performed using Bradford assay (Bio-Rad).

Protein samples were prepared in Laemmli buffer and heated at 95°C for 5 min prior to resolution by sodium dodecyl sulfate-polyacrylamide gel electrophoresis (SDS-PAGE) using 4–15% Mini-PROTEAN TGX Precast Protein Gels (Bio-Rad). Proteins were transferred onto nitrocellulose membranes (Bio-Rad Trans Blot Turbo). Proteins were visualized on membranes using Ponceau S solution (Sigma-Aldrich) prior to 30 min blocking with PBS containing 0.1% Tween (PBS-T) supplemented with 5% milk. Membranes were subsequently incubated with primary antibodies in 5% milk/PBS-T (1:1,000 dilution, except when indicated) for 2h at RT. Primary antibodies used include anti-RIG-I (Alme-1; AdipoGen, 1:1,000), anti-MDA5 (Hely-1; AdipoGen, 1:1000), anti-MAVS (A300-782AM; Bethyl, 1:1,000), anti-IRF3 (D83B9; Cell signaling, 1:1,000) and anti-GAPDH (60004-1-Ig; Proteintech Europe, 1:5,000). Membranes were incubated with horseradish peroxidase (HRP)-coupled secondary antibodies (Cell Signaling Technology) at 1:2,000 dilution for 1h at RT. Immunoreactivity was detected by chemiluminescence (SuperSignal West Pico or Femto, Thermo Fisher Scientific). Images were acquired on an iBright (Invivogen).

### Infectious particles titration

Viral infectivity was quantified by determining the 50% tissue culture infectious dose of 50 per mL (TCID50/mL). Briefly, viral suspensions were serially diluted (10^−1^ to 10^−12^) in cell culture medium and inoculated onto A549 cells seeded in 96-well plates 24 h earlier. After 72 hours, cells were fixed in 2% paraformaldehyde in PBS for 15 min at room temperature, stained with a 2.3% crystal violet in 20% ethanol for 10 minutes, and scored for cytopathic effect. Viral titers were calculated using the Spearman-Karber method.

### RNA extraction and gene expression analyses

Total RNA was isolated with TRIzol reagent (Invitrogen) or Power SYBR Green Cell-to-Ct(Invitrogen). For TRIzol-based RNA extraction, RNA concentration was measured with a Nanodrop spectrophotometer (ND-1000, Nanodrop Technologies), prior to treatment with DNase and cDNA synthesis from 1-2 μg RNA using Owi Script Reverse Transcriptase UltraMix (OZYME) and quantification with a LightCycler 480 cycler (Roche) using Owi Green FAST qPCR Premix (OZYME) and appropriate primers. For Power SYBR Green Cell-to-Ct kit the RT and qPCR were performed following the manufacturer’s instructions using a CFX Opus 384 Real-Time PCR System (Bio-Rad). Relative quantities of the transcripts were calculated using the ΔΔCt method, using the glyceraldehyde-3-phosphate dehydrogenase (*GAPDH*) and ribosomal *18S* RNA for normalization. The RT-qPCR primers used are listed in Table 2,

**Table 2.**
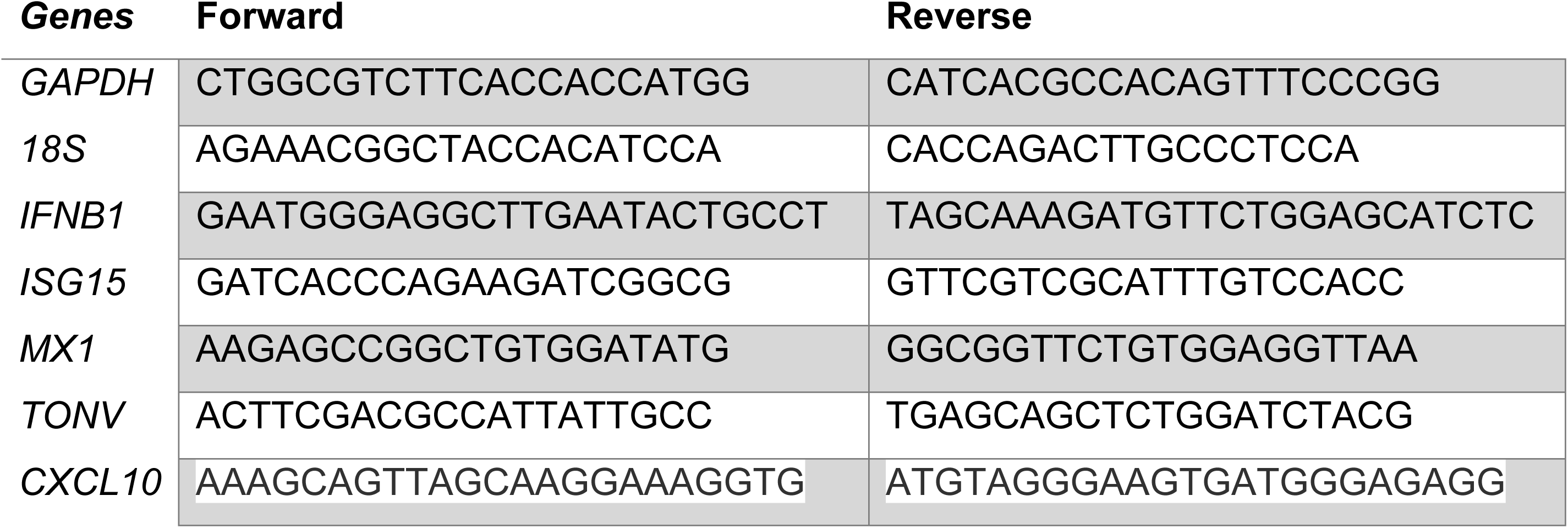
RT-qPCR primers.

### Fluorescence microscopy of T98G upon TONV infection

WT or KO T98G cells were cultured on glass coverslips and infected with TONV at different MOIs. At 24 hours post infection, cell monolayers were rinsed twice with PBS, fixed in PBS-PFA (4%) for 10 min, permeabilized in PBS with triton X-100 (0.05%) for 5 min, and incubated in PBS-BSA (1%) for 30 min. Cells were then labelled with primary anti-dsDNA J2 antibody (Jena Bioscience, RNT-SCI-10010200) at a final concentration of 3 ng/µL in PBS-BSA (1%) incubated overnight at 4°C. The next day, cells were rinsed three times with PBS, and labelled with Alexa fluor A488 (A-11001, Thermofisher scientific) for 2 hours at room temperature. Nuclei were labelled with a PBS-Hoechst solution at 2 μg/mL for 5 min at RT and samples were covered in mounting medium (ProLong P36930, Thermofisher scientific). Labelled cells were analysed with an inverted LSM 880 confocal microscope for qualitative analysis of J2 distribution (Carl Zeiss Ltd.). For quantitative analysis, cells were grown in 96-well plates, followed by the same infection and labelling protocol. Acquisition and analysis of J2 positive cells were performed using an ImageXpress Pico Automated Cell Imaging system.

### IFN-α/β cytokine release assay

Dosage of type I interferons cytokine released during Tonate virus infection was performed using IFN-α/β Reporter HEK 293 Cells (HEK-Blue™ IFN-α/β Cells, InvivoGen). Briefly, 20 µL from the supernatant of T98G infected cells was collected at time points of interest, and transferred into a 96-well culture plate, with the addition of 50,000 cells / well of IFN-α/β Reporter HEK 293 Cells diluted in 180 µL Dulbecco’s Modified Eagle Medium (DMEM) supplemented with 10 % Foetal Bovine Serum (FBS, Eurobio), 1% L-glutamine (Lonza), 1% Penicillin/Streptomycin (Lonza). Reporter cells were maintained overnight in culture at 37°C in presence of test supernatants, before performing type-I IFN dosage assay using QUANTI-Blue™ solution (InvivoGen) according to the manufacturer instructions.

## Data and statistical analysis

Data analysis and plotting were performed using custom Python scripts. All experiments were performed at least three times, and *p*-values were calculated with a student t-test using the open-source library Scipy (scipy.stats.ttest_ind). All data and used python scripts are available upon request.

## Author contributions

Conceptualization, TL, RE, NL, KM; investigation, TL, RE, NL, KM; resources, MM, JR, ZD, MS, MC, IMC, DM, NL,KM. funding acquisition, DM NL, KM.; supervision, DM,NL, KM; writing—original draft, TL, RE, NL, KM; and writing—review and editing, TL, RE, NL, DM, KM.

## Acknowledgments

We thank Dr. Caroline Goujon for providing A549 and T98G cell lines. We thank the CEMIPAI (UAR 3725 CNRS/Montpellier University) BSL-3 Facility for access to their facilities and particularly thank Christine Chable-Bessia and CEMIPAI personnel. We also thank the imaging facility MRI, a member of the national infrastructure France-BioImaging supported by the French National Research Agency (ANR-10-INBS-04). We thank members of the Laguette and Majzoub labs for helpful discussion.

---

**Figure S1.**
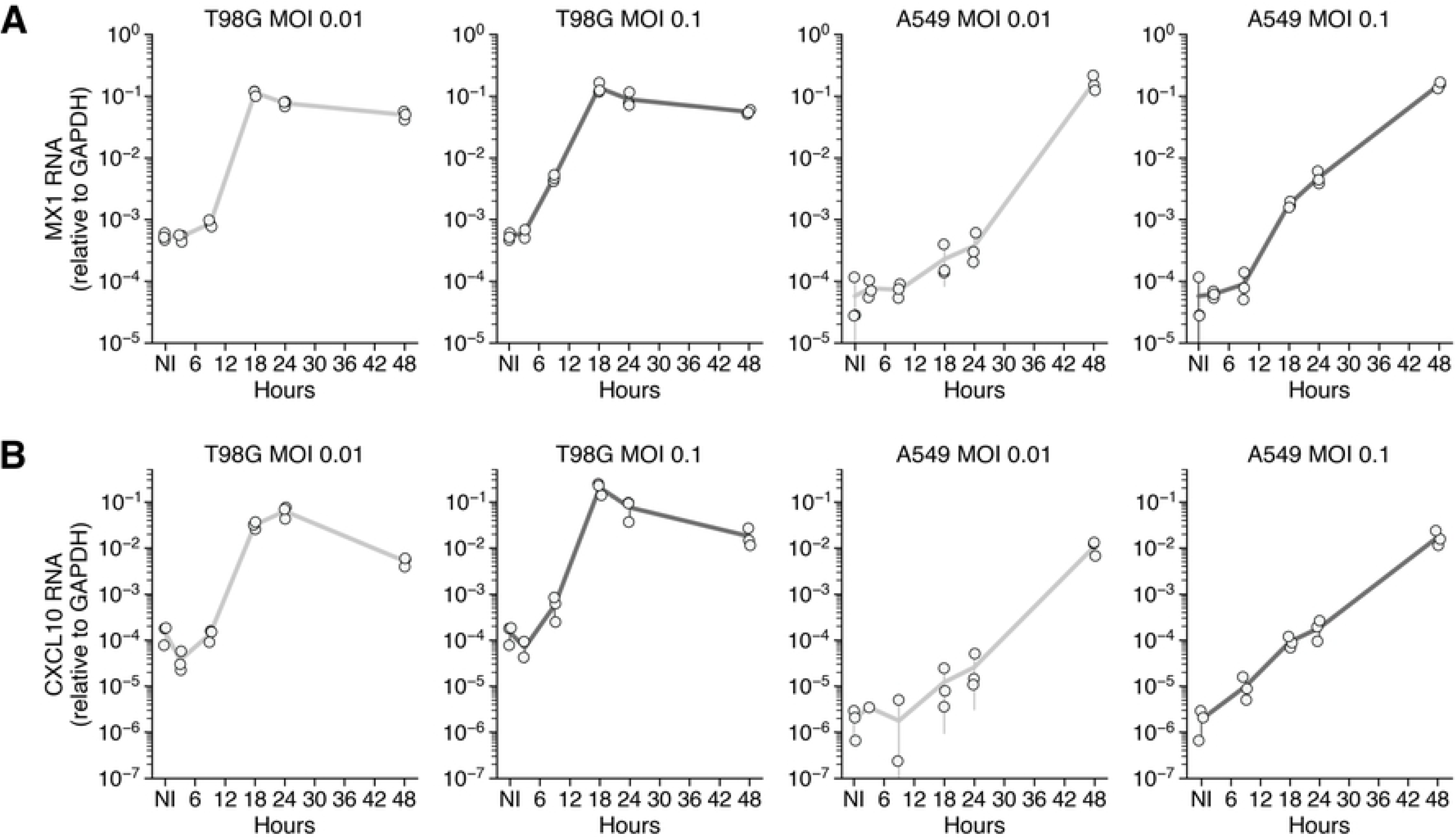

**Figure S2.**
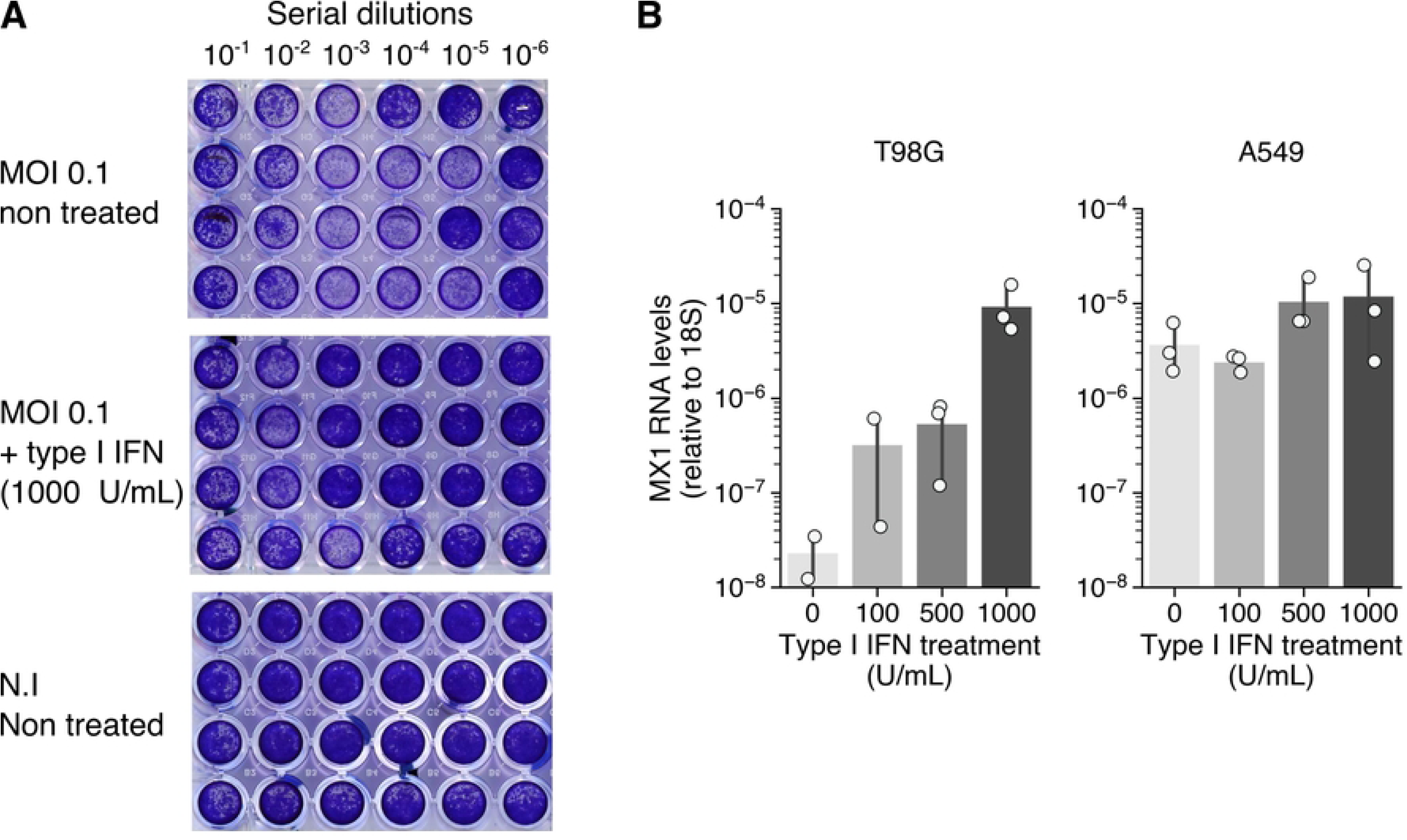

**Figure S3.**
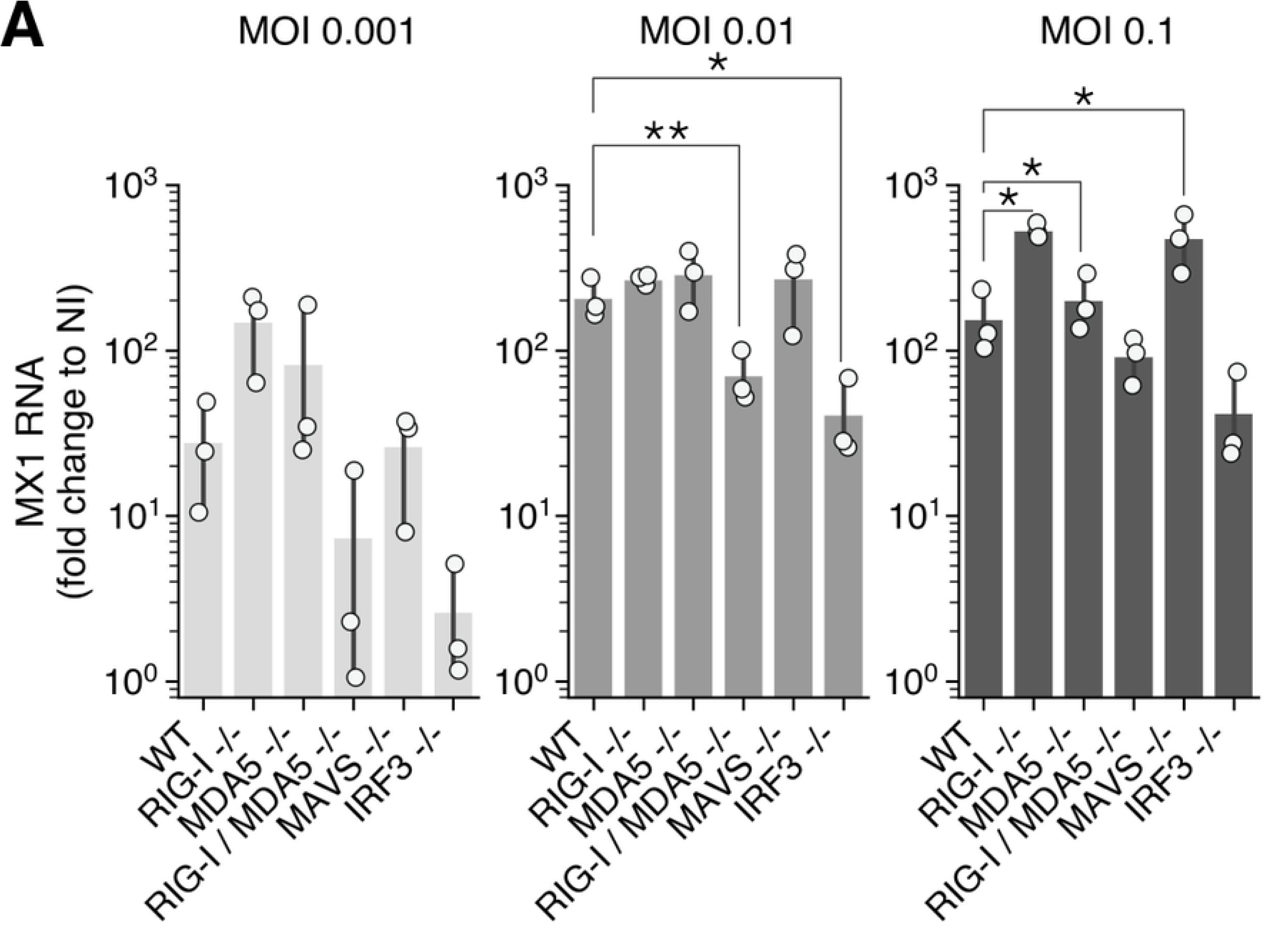

